# ForageGrassBase: Molecular resource for the forage grass *Festuca pratensis* Huds

**DOI:** 10.1101/674390

**Authors:** Jeevan Karloss Antony Samy, Odd Arne Rognli, Mallikarjuna Rao Kovi

## Abstract

**Background:** Meadow fescue (*Festuca pratensis* Huds.) is one of the most important forage grasses in temperate regions. *F. pratensis* is a diploid (2n =14) outbreeding species that belongs to the genus Festuca. Together with Lolium, they are the most important genera of forage grasses in temperate regions. *F. pratensis* has good winter survival, with high quality dry matter yields and persistency, and is suitable both for frequent-cutting conservation regimes and for grazing. It is a significant component of species-rich permanent pastures in the temperate regions, ensuring high forage yield under harsh climatic conditions where other productive forage grass species are unable to grow. However, genomic resources for *F. Pratensis* is not available so far.

**Results:** The draft genome sequences of two *F. pratensis* genotypes “HF7/2” and “B14/16” are reported in this study. Here, using the draft genome, functional annotation datasets of two *F. pratensis* cultivars, we have constructed the *F. pratensis* genome database http://foragegrass.org/, the first open-access platform to provide comprehensive genomic resources related to this forage grass species. The current version of this database provides the most up-to-date draft genome sequence along with structural and functional annotations for genes using Genome Browser (GBrowse). In addition, we have integrated comparative genomic tracks for *F. pratensis* genomes by mapping *F.pratensis* genome to the barley, rice, Brachypodium and maize genomes. We have integrated homologus search tool BLAST also for the users to analyze their data. Combined, GBrowse, BLAST and downloadble data gives an user friendly access to *F. pratensis* genomic resouces. All data in the database were manually curated.

**Conclusion:** To our knowledge, ForageGrassBase is the first genome database dedicated to forage grasses. It provides valuable resources for a range of research fields related to *F. pratensis* and other forage crop species, as well as for plant research communities in general. The genome database can be accessed at http://foragegrass.org. In the near future, we will expand the ForageGrassBase by adding genomic tools for other forage grass species, as soon as their genomes become available.

## Background

Grasslands are covering very large portions of the earth’s surface and they are important as feed sources and pastures for livestock. In both developed and developing countries, many millions of livestock farmers, ranchers and pastoralists depend on grasslands and conserved products such as hay and silage from a range of fodder crops for their livelihoods. Among several forage crops, meadow fescue (*Festuca pratensis* Huds.) is one of the most important forage grass species in temperate regions of the world. It is a diploid (2n =14) outbreeding species that belongs to the genus Festuca (1)

Fescues in general have evolved superior adaptations to abiotic stresses, e.g., winter survival in meadow fescue. The most abundant forage grass species in temperate regions, perennial ryegrass (*Lolium perenne* L.), is known for its superior nutritive quality, rapid establishment and growth but is lacking persistency under harsh environmental conditions. The Lolium-Festuca species complex is unique since it is possible to combine Lolium and Festuca genomes in interspecific hybrids (Festulolium) (2). Complementation of traits in Festulolium hybrids is thus a very interesting strategy for developing novel germplasm and cultivars with improved quality and persistency, which can contribute to a sustainable forage production. Relatively modest genomic resources have been developed for meadow fescue compared with other grass species like perennial ryegrass (*Lolium perenne*) (3).

In order to develop better Festulolium hybrids, we have initiated sequencing of *F. pratensis*, and combined with an efficient utilization of the close relationship with barley (*Hordeum vulgare*), rice (*Oryza sativa*), *Brachypodium distachyon* and maize (*Zea mays*) through comparative genomics approaches. High quality annotated Festuca genomes are now available. As a first step, the genome sequences and genome annotations for two *F. pratensis* genotypes are made available through ForageGrassBase (http://foragegrass.org). ForageGrassBase was developed to make these substantial amounts of genomic data accessible through visualizations and analytic tools in a common framework. Similar resources for other forage grass species will be added to ForageGrassBase when they become available.

## Construction and content

Bootstrap (HTML, CSS), Javascript, PHP and Python were used to develop ForageGrassBase. The Generic Genome Browser (GBrowse) [4] and BLAST [5] were also installed. R packages are used for BLAST results visualizations.

*De novo* sequencing of the *Festuca pratensis* genomes were performed using Illumina mate pair sequencing and assembly was performed by the SOAPDenovo assembler. Furthermore, gene annotation was performed by in-house developed annotation pipelines and python scripts.

## Utility and discussion

### Genome browser (GBrowse)

The generic genome browser (GBrowse) is simple and one of the most used genome browsers for visualization of genomes. We installed GBrowse to visualize and share genomic data of *F. pratensis* (Fig. 1). Currently, ForageGrassBase contains molecular data of two *Festuca pratensis* genotypes; Festuca HF2/7, a Norwegian genotype originating from a population selected for high frost tolerance and a Yugoslavian genotype, B14/1700, which is used by our group to develop a mapping family for linkage map construction (6). Further, a comparative genome analysis was performed against other grass species like barley, Brachypodium, rice and maize. These comparative genomics tracks consisting of gene names and chromosome positions were added to the genome browsers (Fig. 1). More data and tracks will be added in the near future for other economically important forage grass species like timothy (*Phleum pratense*) to expand the forage grass genomics resources in ForageGrassBase.

**Fig. 1.**
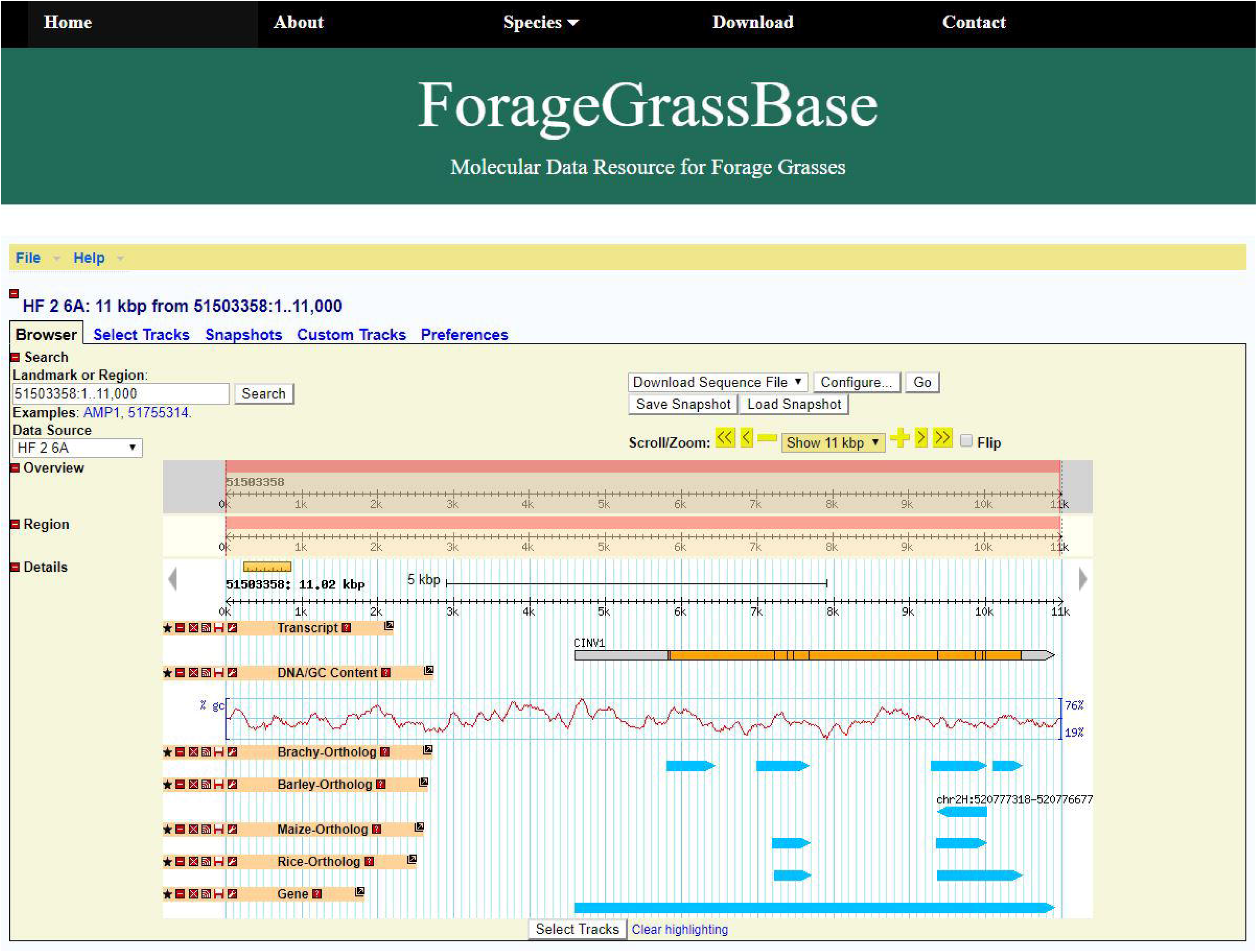
ForageGrassBase genome browser contains *F. pratensis* genome annotation and their orthologous regions in Brachypodium, barley, maize and rice.

### BLAST server

We have installed a BLAST server to search for homologous regions in the *F. pratensis* genome. Users having unknown sequences can use BLAST search to find the homologous regions in Festuca and their corresponding homologous genes and their physical location in Brachypodium,, barley, rice and maize (Fig. 2a). After the search, our algorithm chooses the best hits and plots them in a unique way. BLAST results are connected to GBrowse, so the users can view the homologous regions and nearby genes and other genomic features in all these species (Fig. 2b).

**Fig. 2.**
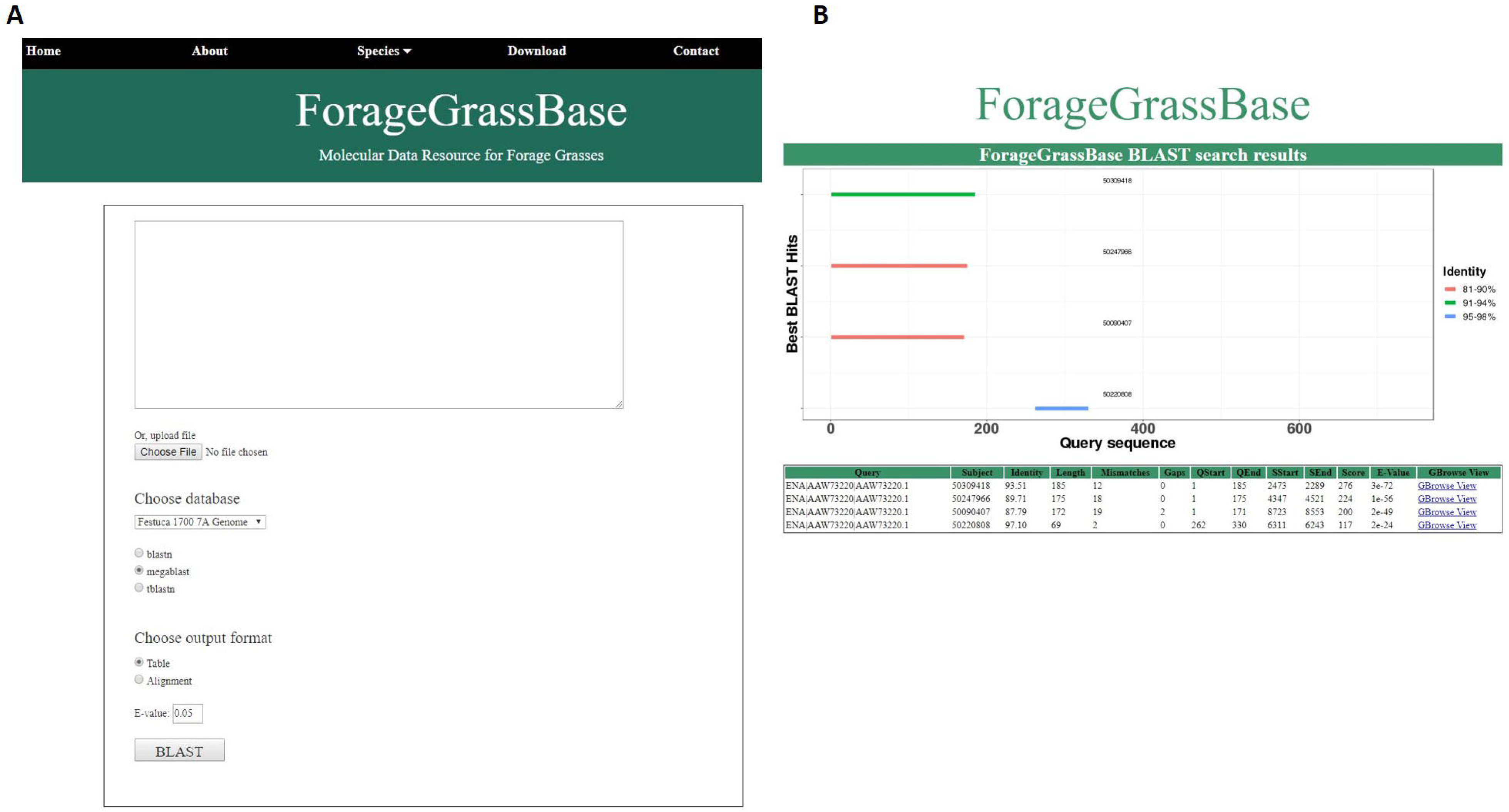
*F. pratensis* cultivars genome browsers with genome annotations and BLAST tool (a) BLAST tool implemented to search for homologous regions in the reference genomes available. (b) BLAST results page shows the homologous regions.

### Future plans and integrations

ForageGrassBase was developed based on high interest for the molecular data of *Festuca pratensis*. Genetic variations and gene expression data will be added using Genetic variation browser (GVBrowser) and Gene expression browser (GEBrowser) in the very near future. Due to rapid developments and lower costs of high-throughput sequencing technologies, we expect more forage grass genome sequence data to be available soon, and these resources and new tools will be added under ForageGrassBase.

### Database access and feedback

All the data used in developing this database are available through the ‘Download’ menu in ForageGrassBase. Genome sequences and gene annotation files for the two Festuca genotypes are available in “fasta” and “gff3” file formats to download and re-use. Users can send their questions and comments through ‘Contact form’ under ‘Contact’ menu.

## Conclusions

To the best of our knowledge, ForageGrassBase is the only online database to access, visualize and download data for the forage grass species *Festuca pratensis* and its homologous sequences/genes in rice, barley, Brachypodium and maize. Due to rapid developments in high-throughput sequencing technology in recent years, we expect a huge influx of data for forage grasses. Thus, the database ForageGrassBase will be updated by adding new forage grass species genomic resources as soon as their genome sequences are publicly available.

## List of abbreviations

*F. pratensis*: *Festuca pratensis*
GBrowser: Genome browser
BLAST: Basic Local Alignment Search Tool

## Declarations

## Acknowledgements

We thank Torben Asp, Aarhus University, Dag Inge Våge, Teshome Dagne Mulugeta, Torfinn Nome and the Orion computational facility at the Norwegian University of Life Sciences for their support.

## Funding

This project has received financial support from the Research Council of Norway, project numbers: 199664/I10 (VARCLIM), 255428 (GenSelTim) and 208481 (ELIXIR.NO). The funding source had no role in study design, data collection and interpretation and in writing the manuscript.

## Availability and requirements

ForageGrassBase can be accessed at http://foragegrass.org/

## Availability of data and materials

This work does not contain additional data.

## Authors’ contributions

MRK, JKAS and OAR conceived the idea of developing ForageGrassBase. OAR, MRK provided the genome sequences and annotation files. JKAS developed ForageGrassBase with inputs from MRK and OAR. JKAS and MRK wrote the manuscript and included comments from OAR. All authors read and approved the final manuscript.

## Competing interests

The authors declare that they have no competing interests.

## Consent to publish

Not applicable.

## Ethics approval and consent to participate

Not applicable

## Notes

https://foragegrass.org/

